## Supplementary_figures for "Sunpheno: a deep neural network for phenological classification of sunflower images"

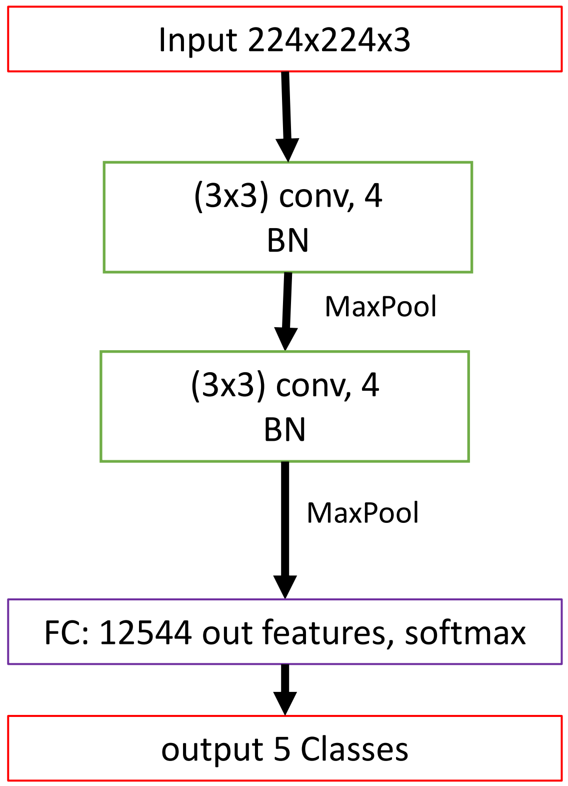


**Figure supplementary 1:** Architecture of CNN from scratch method. The input is an image with dimensions of height (224 px), wide (224 px) and three channels RGB (224x224x3). After passing through two convolution layers a fully connected (FC) layer described the image in 12544 features, those features are used to classify the input into 5 classes. Convolution layers (Conv), batch normalizations (BN), karnel size it is shown between brackets.

**Figure supplementary 2:** Architecture of Vgg19 methods. The input is an image with dimensions of height (224 px), wide (224 px) and three channels RGB (224x224x3). After passing through the layers a fully connected (FC) layer described the image in 4096 features, those features are used to classify the input into 5 classes. Convolution layers (Conv), batch normalizations (BN), Relu activation function, karnel size it is shown between brackets.

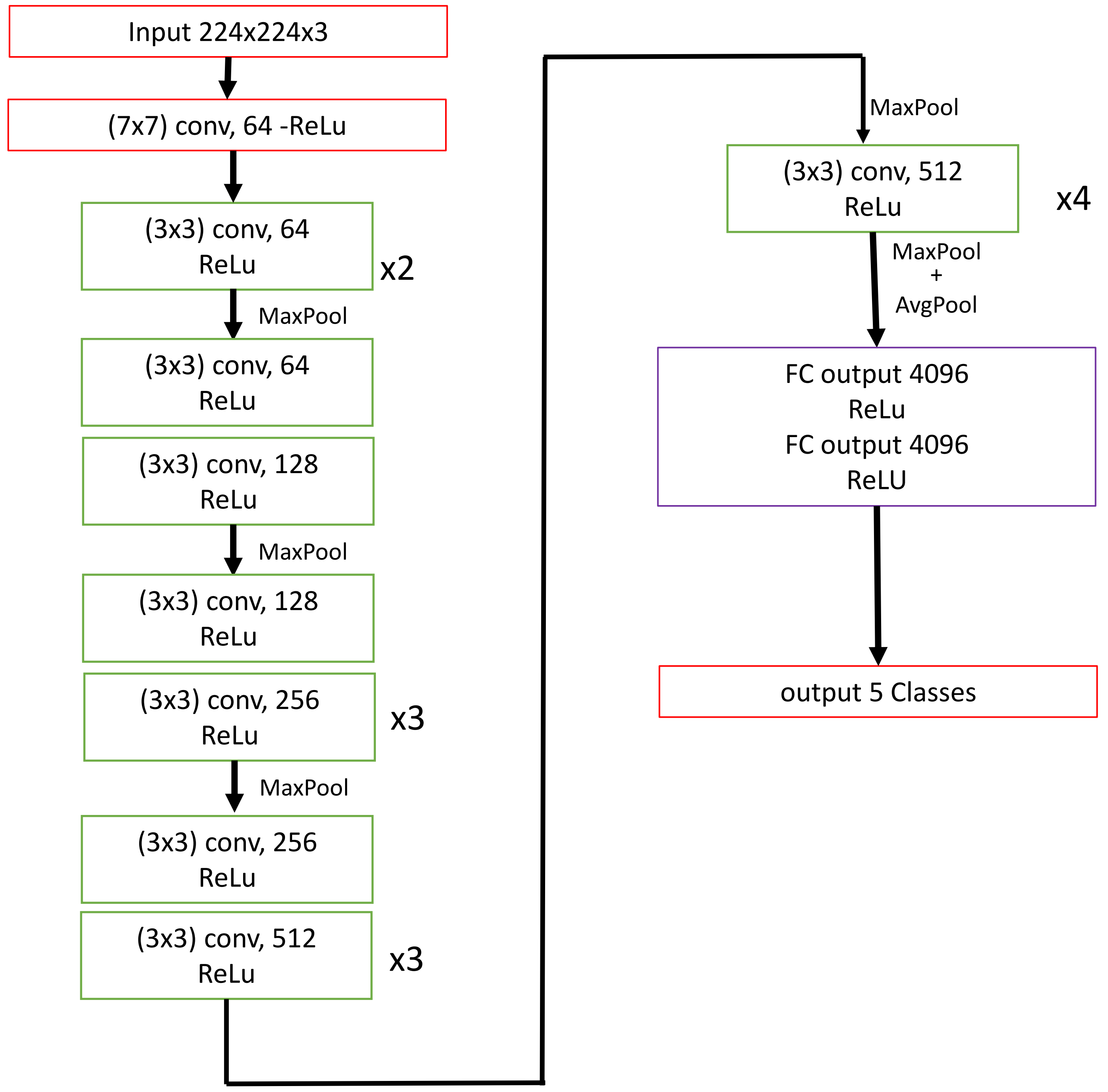

**Figure supplementary 3:** Architecture of ResNet methods. (a) ResNet-18 (b) ResNet-50. The input is an image with dimensions of height (224 px), wide (224 px) and three channels RGB (224x224x3). After passing through the layer1, layer2. layer3, layer4  a fully connected (FC) layer described the image in 1000 features, those features are used to classify the input into 5 classes. Each layer (1-4) consists repetition of residual building block (RBB) each RBB has convolution layers (Conv), batch normalizations (BN), Relu activation function. The symbol (*) is used to imply that at the last RBB of the layer contains an extra Conv and BN layers. Karnel size it is shown between brackets.

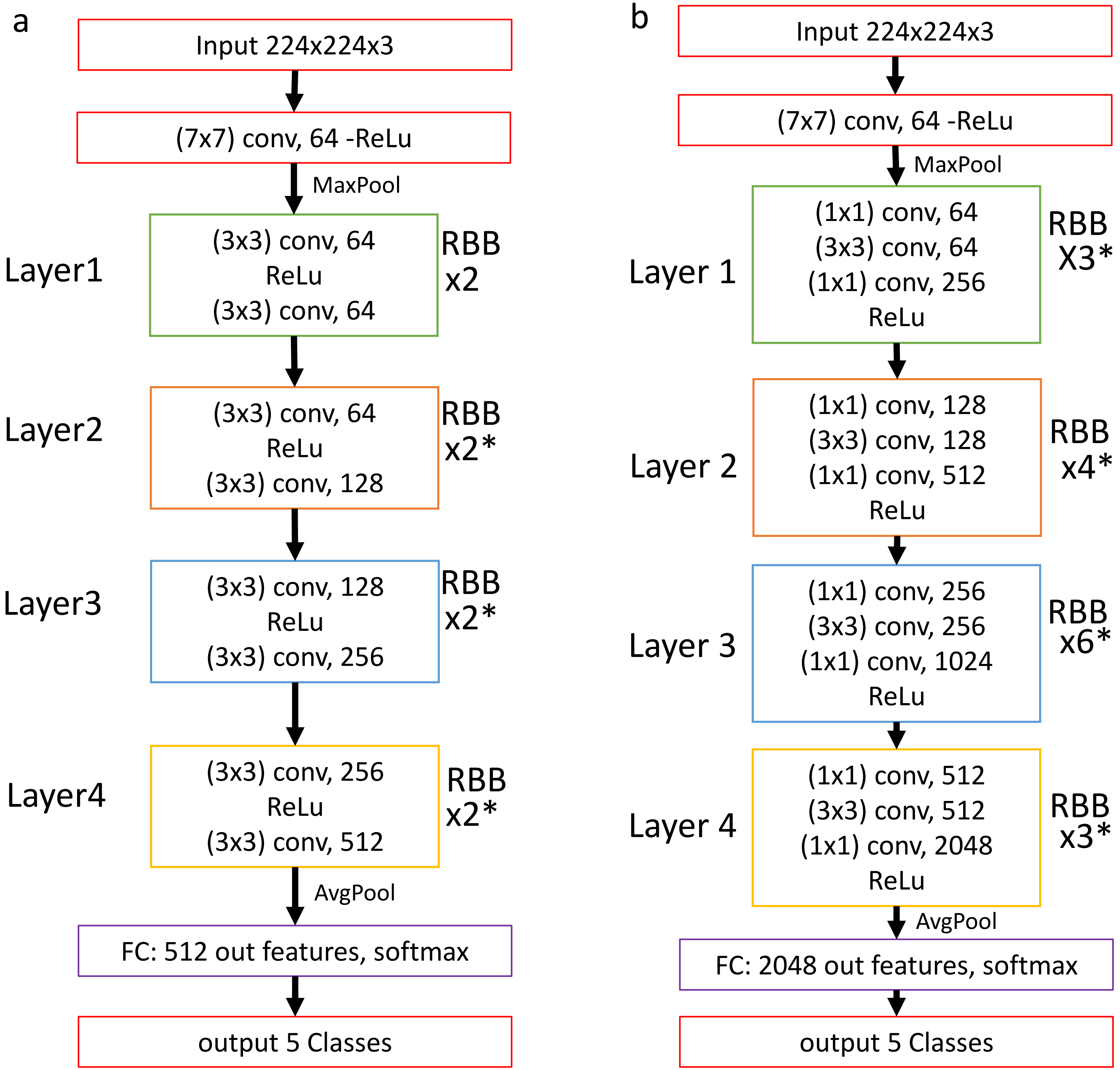

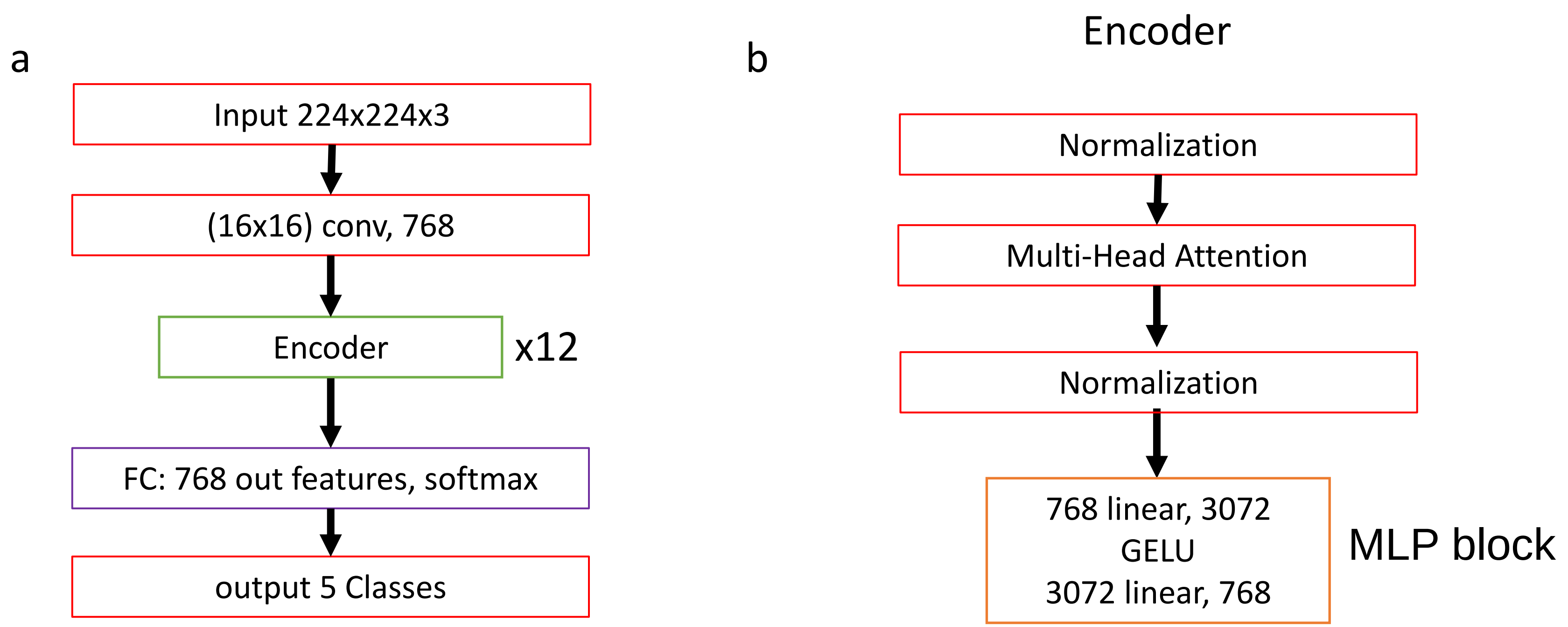


**Figure supplementary 4:** Architecture of Vit-B16 methods. (a) The input is an image with dimensions of height (224 px), wide (224 px) and three channels RGB (224x224x3). After passing through a convolution layer, 12 encoder layers a fully connected (FC) layer described the image in 768 features, those features are used to classify the input into 5 classes. (b) each decoder layer contains a Multi-Head Attention and a multi-layer perceptron (MLP block). Convolution layers (Conv), Gelu activation function, karnel size it is shown between brackets.
